# The impact of clade B lineage 5 MERS coronaviruses spike mutations from 2015 to 2023 on virus entry and replication competence

**DOI:** 10.1101/2025.06.30.662263

**Authors:** Ray TY So, Kaman KM Lau, Ziqi Zhou, Leo LM Poon, Malik Peiris

## Abstract

Middle East respiratory syndrome coronavirus (MERS-CoV) is an emerging coronavirus that can cause zoonotic disease in humans with lethal severe viral pneumonia. Dromedary camels are the source of zoonotic infection. As of June 2025, MERS-CoV has resulted in a total of 2626 reported cases, 36% of these being fatal. The number of reported human cases has been on a decreasing trend since 2016 and reached a minimum level during the COVID-19 pandemic. The reason for the reduction of cases is unclear and may be multifactorial. We hypothesized that mutations accumulating in the virus spike protein may have reduced zoonotic potential. Here, we investigate the impact of recently emerged virus spike-protein mutations on virus replication competence using pseudoviruses and replication-competent recombinant viruses. We found that two spike variants detected in 2019 show a reduced cell entry and lower viral replication in human cells. However, spike variants detected in 2023 sequences, did not show significant changes in cell entry and viral replication. All the MERS-CoV spikes tested showed a cell-entry pathway preference via the cell-surface TMPRSS2 route. Our data suggests that spike protein mutations are not a major determinant of the fewer MERS-CoV human cases observed.

**Author Summary:** MERS-CoV is identified by the World Health Organization (WHO) as a potential pandemic candidate. The ability of coronaviruses to mutate and adapt in new hosts raises concerns about the impact of virus genetic changes on human zoonotic potential. There has been a notable decline in human MERS cases reported to the WHO since 2018, but the underlying reasons remain unclear. Here, we focus on investigating whether the recently emerged virus spike mutations may contribute to this observation. We found that while some spike mutations detected in 2019 reduce cell entry and viral replication, more recent viruses do not share this phenotype. This study highlighted a need for comprehensive genomic surveillance and phenotyping of recent MERS-CoV isolates to understand the potential role, if any, of other non-spike virus mutations on viral zoonotic competence and to explore alternate hypothesis, such as cross-reactive immunity from COVID-19 contributing to reduced human MERS-CoV disease.

## Introduction

Middle East respiratory syndrome coronavirus (MERS-CoV) is a highly pathogenic zoonotic coronavirus that was first reported in a patient with severe viral pneumonia in Saudi Arabia in 2012 [1]. Dromedary camels are recognized as a major source of zoonotic infection [2]. Surveillance studies subsequently revealed MERS-CoV circulation in dromedary camels in the Arabian Peninsula, Northern, Eastern and Western Africa, and in the Central Asia, e.g. Pakistan [3-11]. While camel exposure was associated with an increased risk of human disease, the exact camel-to-human transmission mechanisms remain unclear. As of June 2025, MERS-CoV has resulted in a total of 2626 reported human cases, causing 947 deaths with a fatality rate of 36% (WHO statistics [12]). These reported cases include primary zoonotic infections, as well as clusters of human-to-human transmissions in hospital settings, as documented in Saudi Arabia [13, 14]. In 2015, a single returning traveler from the Arabian Peninsula to the Republic of South Korea initiated an outbreak resulting in a total of 186 confirmed cases [15]. Increased awareness and improved hospital infection prevention and control measures have contributed to the reduction of MERS cases and deaths since 2016 [16]. Interestingly, since 2017, there was a further reduction of human cases reported from the Arabian Peninsula (WHO statistics, Fig 1A). The trend of reduced human cases has persisted even after the relaxation of public health and social restriction measures implemented during the COVID-19 pandemic. The reason for the reduced case reports of MERS may be multifactorial, including better infection control measures in hospitals, increased public awareness to coronavirus diseases, cross-protection from COVID vaccination or infection, or changes in the MERS-CoV phenotype. In this study, we investigate the potential impact of recently emerged mutations in the virus spike protein on virus replication competence, as one potential explanation.

**Figure 1.**
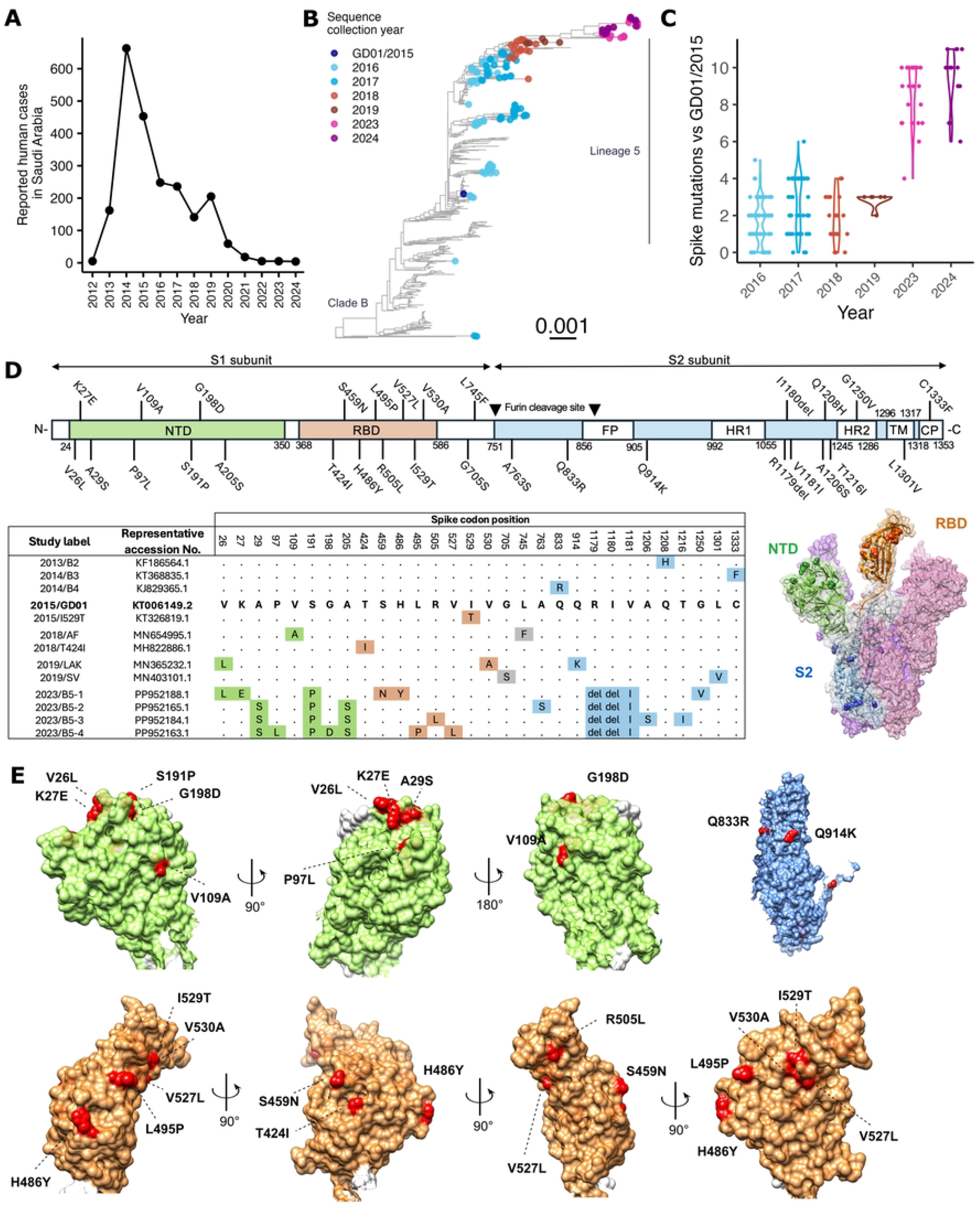
MERS-CoV human cases and the accumulation of spike amino acid mutations. (A) Numbers of lab confirmed human MERS-CoV cases in Saudi Arabia reported to the World Health Organization (WHO) [12]. (B) Full genome ML tree of 607 MERS-CoV sequences. Reference sequence MERS-CoV/GD01 collected in 2015 and sequences with collection year in 2016 and later are colored. Scale bar represent nucleotide change per site. (C) Dot and violin plot of number of spike amino acid mutation from sequences collected from 2016 to 2024 against the reference sequence 2015/GD01. (D) Illustration showing the domains of MERS-CoV spike protein and the position of spike mutations tested in this study. Study labels and representative accession number of sequences carrying the indicated spike mutations are listed. Mutations are located on the spike protein structure based on the PDB:5×59 file. Domains are color for NTD as green, RBD as orange and S2 as blue. (E) Mutations on the spike surface for each domain are highlighted as red using Chimera software.

The genetic evolution of MERS-CoV primarily occurs within the camel hosts [17]. Multiple lineages of MERS-CoVs have been shown to circulate among camels, which later underwent genetic recombination to result in a recombinant lineage (clade B lineage 5), which was associated with large outbreaks in 2015 [6]. Experimental data showed the clade B lineage 5 had a higher viral replication fitness in vitro and ex vivo compared with earlier strains [18]. A recent study showed MERS-CoV sequences collected from camels in Saudi Arabia in 2023-2024 had emerged from previous clade B lineage 5 strains and had acquired unique genetic features [19]. Spike mutations in MERS-CoV have been shown to modulate viral replication in-vitro and pathogenicity in a hDPP4-knockin mouse model [20]. The spike I529T mutation has been reported to have a lower cell entry, reduced viral replication and an attenuated phenotype in the mouse model [20, 21]. Similarly, spike mutations in an African clade C strain from Burkina Faso have also demonstrated reduced viral production in human respiratory tract tissues [22]. These observations suggest that MERS-CoV spike protein can be one major molecular determinant of the virus phenotype in humans.

Here, we hypothesised that mutations in spike protein in more recent clade B MERS-CoV may contribute to reduced pathogenic potential to humans, resulting in reduced numbers of zoonotic MERS in recent years. We analysed MERS-CoV spike protein mutations observed since 2015 and generated spike-packaged pseudoviruses to assess any change of cell entry from representative spike proteins. We also developed a Golden Gate assembly reaction that allowed efficient generations of full-length MERS-CoV encoding pBAC infectious clones with different chimeric spike protein mutants on the clade B lineage 5 2015/GD01 MERS-CoV backbone. By comparing the replication kinetics among these recombinant chimeric spike MERS-CoVs on the same genetic backbone, the phenotypic effect contributed by the spike protein can be assessed. Mechanistically, we also investigated potential changes in the entry pathway preference contributed by the spike mutations. Compared with clade B lineage 5 2015/GD01 virus as reference, our data showed spike mutants from 2019 had reduced human cell entry and viral replication, but virus spikes from 2023 viruses did not demonstrate significant differences. Our findings suggest that mutations in the virus spike protein do impact on viral replication phenotype (e.g. in 2019), but are not a likely cause for the fewer human MERS cases observed in the Arabian Peninsula in more recent years.

## Results

### Spike mutations accumulate during MERS-CoV evolution

We retrieved human and camel MERS-CoV sequences available from Genbank and included newly published sequences collected from camels in Saudi Arabia in 2023-2024 for our analysis (total n = 607) [19]. Phylogenetic analysis showed MERS-CoVs in 2016-2017 had diverged into multiple clade 5 sub-lineages in 2016-2017, but a single sub-lineage dominated from 2018 onwards (Fig1B). Recent virus sequences from 2023 and 2024 were monophyletically clustered with those from 2019, suggesting that these recent viruses shared a recent common ancestor and a direct evolutionary link. MERS-CoV spike protein is responsible for host cell entry and determines the host-species tropism. Given the limited number of human cases reported after COVID-19, we aimed to investigate whether the spike protein plays a major role explaining the recent changes in epidemiology of MERS-CoV in the Arabian Peninsula. We used the ancestral clade B lineage 5 2015/GD01 strain, a virus representing the 2015 outbreak in South Korea, as the reference of this study. By aligning the spike protein sequences, we observed an average number of 1.7 to 2.9 amino acid substitutions in viruses from 2016 to 2019 sequences, respectively (Fig1C). The number of amino acid substitutions further increased in 2023 and 2024 sequences, with an average of 8.7 and 9.4 mutations, respectively, relative to 2015/GD01.

The observed spike mutations occurred in spike S1 N-terminal domain (NTD), receptor binding domain (RBD) and spike S2 domains (Fig1D). Structural analysis revealed most of these mutations NTD were located at the surface of the NTD, with the exception of the A205S mutation (Fig1E). Similarly, all eight mutations on the RBD were located on the protein surface. The Q833R and Q914K mutation were located the surface of the S2 trimer. The lack of sequence data from 2020-2022 hindered a more sequential tracking of how spike mutations in 2023 emerged.

To investigate the phenotypic effect of these observed mutations, we picked representative sequences from early clade B lineages (2013/B2, 2014/B3 and 2014/B4), 2018 (2018/AF and 2018/T424I), 2019 (2019/LAK and 2019/SV) and 2023 (2023/B5-1, 2023/B5-2, 2023/B5-3 and 2023/B5-4), to compare their cell-entry using pseudotyped viruses expressing representative spike proteins and the effect on viral replication in-vitro using infectious chimeric 2015/GD01 viruses expressing the selected virus spike proteins. The I529T mutation previously identified in 2015 Korean sequences associated with reduced human DPP4 binding and mice pathogenicity was used as an additional control [20, 21].

### Impact of MERS-CoV spike mutations on cell-entry

We generated lentiviral-pseudoviruses expressing different spike proteins and measured cell-entry in Vero and Calu3 cells. It has been shown that MERS-CoV utilizes the endocytic pathway for entry into Vero cells, whereas virus entry into Calu3 cells are mediated by transmembrane protease, serine 2 (TMPRSS2) activation at the cell surface [23]. We first tested early clade B lineage 2, 3 and 4 spike proteins, which contain mainly S2 Q1208H, C1333F and Q833R mutation respectively, and showed they did not show significant changes in cell-entry relative to 2015/GD01 spike (Fig 2A). Similarly, 2018/AF, which contains V109A and L745F mutations, and 2018/T424I, with an T424I mutation at the RBD, also showed no significant difference in virus entry compared with 2015/GD01. Next, we tested spike from 2019 sequences and observed significant reduction of cell-entry in both Vero and Calu3 cells for 2019/LAK and 2019/SV spike (Fig 2B), the former showing greater reduction compared to 2019/SV in both cell types. The extent of reduction of cell entry of 2019/LAK was comparable to the previously reported virus mutation 2015/I529T. Interestingly, the 2019/LAK virus contains a V530A mutation, which is located adjacent to the I529T mutation. We further identified that the V530A mutation in 2019/LAK and the V1301L mutation in 2019/SV were the main determinant of the cell entry in both spikes (Fig 2C). The V1301L mutation with a reduced cell entry is located at the transmembrane domain of the spike protein. We did not observe any reduction in cell entry associated with the 2023 sequences, although 2023/B5-3 showed a weak but significant reduction of cell entry of 0.23 log_10_ RLU (Fig 2D).

**Figure 2.**
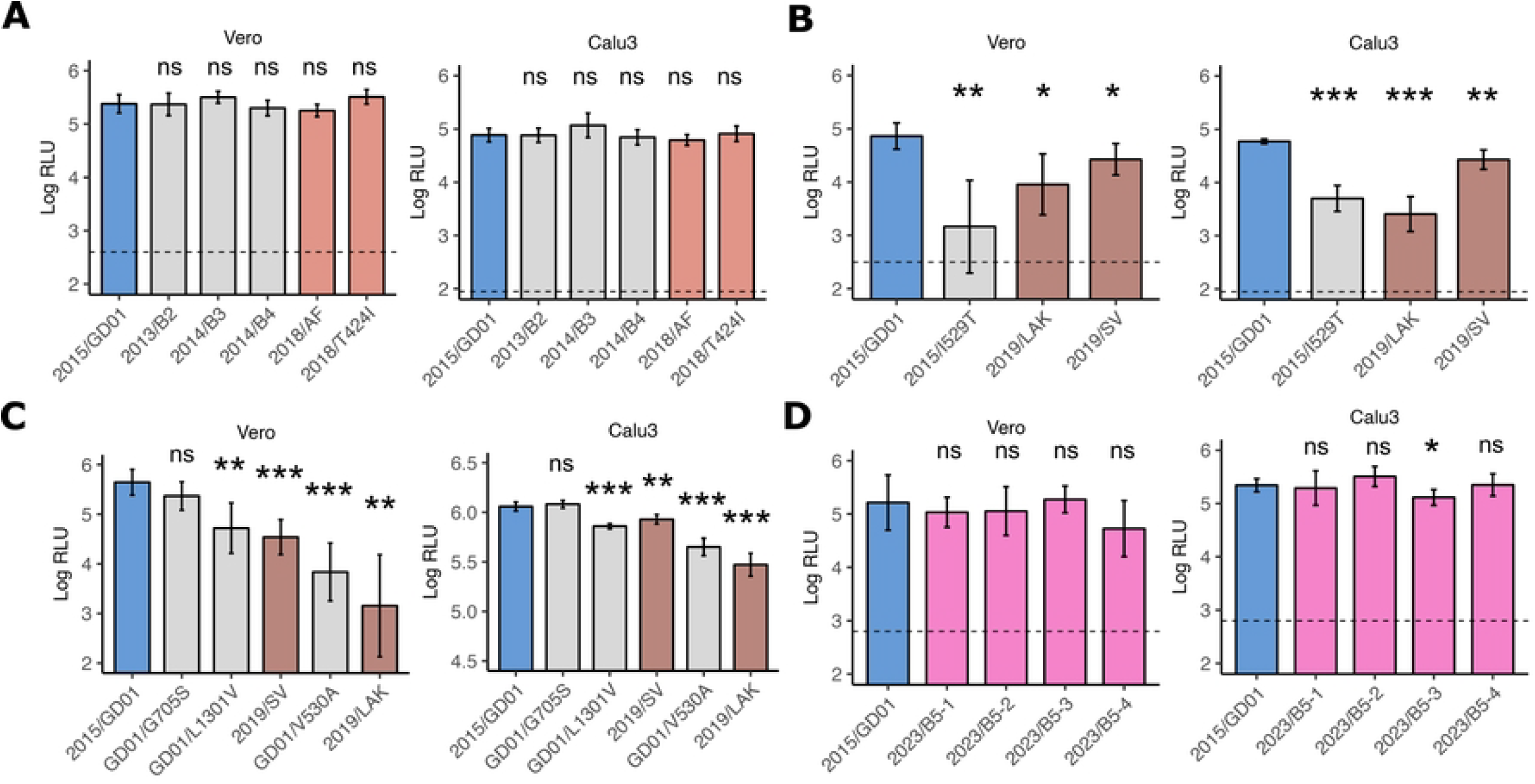
Cell entry of spike mutations using pseudovirus assay. (A-D) Cell entry of spike protein expressed lentivector pseudoviruses in Vero and Calu3 cells were measured by relative luciferase units (RLU). Spike expression plasmids encoding (A) clade B lineage 5 viruses from 2015 to 2018, (B) sequences from 2019, (C) selected single mutations from 2019 viruses, and (D) sequences from 2023, were co-transfected with third-generation lentivirus expression plasmids in 293T cells. In (A)-(D), a normalized dose of 7-8 log_10_ p24 RNA copies per 100ul of pseudovirus supernatant was added to target cells. Dotted horizontal line showed the RLU from “bald” non-spike expressed pseudovirus controls. Assays were performed with five replicates using two independent batch of pseudoviruses (A to D) and the data are presented as the average ± SD. Statistically significant differences between 2015/GD01 and mutant spikes pseudoviruses were determined by two-sided Student’s t tests (* p < 0.05, ** p < 0.01, *** p < 0.001).

### In vitro replication kinetics of recombinant viruses with representative spike mutants in comparison to MERS-CoV 2015/GD01

In order to confirm the data obtained from pseudotyped viruses, we sought to assess the impact of selected virus spike mutations in infectious chimeric 2015/GD01 MERS-CoV expressing the mutant virus spikes. A detailed bio-risk analysis was carried out prior to the initiation of these experiments (see methods). As the spike mutations introduced to the chimeric viruses were all reported to have naturally occurred in the field, these studies were not considered “gain-of function” research. All studies with recombinant virus rescue and characterization were carried out in biosafety-level-3 containment.

We utilized the Golden Gate assembly strategy to modify the pBAC infectious clone encoding a MERS-CoV 2015/GD01 genome into different pBAC infectious clones encoding chimeric recombinant MERS-CoV 2015/GD01 backbone with different spike variants (Fig. 3A). First, fragments of 2015/GD01 genome were cloned into 10 individual high-copy pUC vector plasmids. Plasmid pUC-F8, encoding the spike gene, was then induced with the mutations using site-mutagenesis or direct gene synthesis. Plasmids pUC-F1 to F10, together with a plasmid encoding the pBAC backbone, were pooled together in a Golden Gate assembly reaction to generate the circular pBAC infectious clone encoding different chimeric spike recombinant MERS-CoV 2015/GD01 viruses (Fig3B and 3C). A total of 7 recombinant viruses were rescued and tested in this study: 2015/GD01 (reference virus), 2015/I529T, 2018/T424I, 2019/LAK, 2019/SV, 2023/B5-3 and 2023/B5-4.

**Figure 3.**
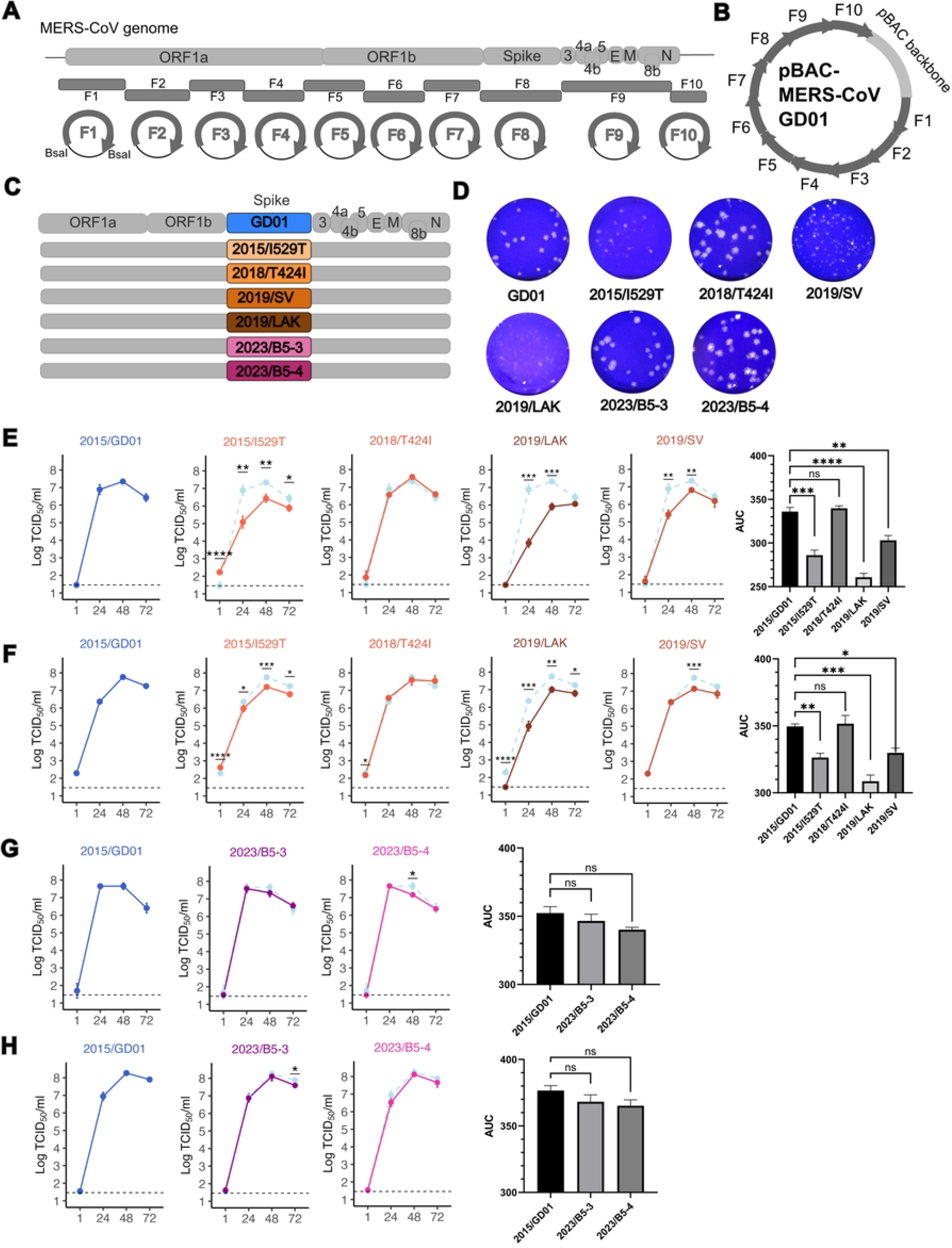
Replication kinetics of recombinant MERS-CoV/GD01 with chimeric spike variants. (A) Scheme of the strategy to generate infectious clone encoding recombinant MERS-CoV/GD01 with chimeric spike variants using the Golden Gate assembly. (B) A 38kb circular pBAC infectious clone encoding the complete genome of a recombinant MERS-CoV 2015/GD01. (C) MERS-CoV genome and its individual genes. Spike genes encoding the mutant variants were swapped with the original GD01 spike. (D) Plaque assay of rescued recombinant MERS-CoV/GD01 viruses in Vero cells. (E-F) Replication kinetics of recombinant MERS-CoV/GD01 encoding 2015/I529T, 2018/T424I, 2019/SV and 2019/LAK vs 2015/GD01 in (E) VeroE6/TMPRSS2 cells and (F) Calu3 cells. (G-H) Replication kinetics of recombinant MERS-CoV/GD01 encoding 2023/B5-3 and 2023/B5-4 vs 2015/GD01 in (G) VeroE6/TMPRSS2 cells and (H) Calu3 cells. (E-H) Horizontal dotted line in the line chart represents the limit of detection for the TCID_50_ assay. Figures show the representative data from 2 (G-H) to 3 (E-F) independent experiments each with three biological replicates. Each dot represents the average ± SD. The bar chart represents the AUC analysis using datapoints from 24, 48 and 72 hpi. Error bars indicate the standard error of the mean. Statistically significant differences between 2015/GD01 and other spikes were determined by two-sided Student’s t tests (* p < 0.05, ** p < 0.01, *** p < 0.001).

We observed that 2015/I529T, 2019/LAK and 2019/SV showed smaller plaque size in Vero cells relative to 2015/GD01, but 2018/T424I, 2023/B5-3 and 2023/B5-4 viruses showed no significant difference (Fig3D). As expected, 2015/I529T demonstrated similar smaller plaque morphology compared with 2015/GD01, as previously reported by other studies using a clade A recombinant MERS-CoV/EMC backbone [20, 24].

We compared the growth kinetics of these viruses in VeroE6-TMPRSS2 and Calu3 cells by infecting cells at a low multiplicity-of-infection (MOI) of 0.01. In VeroE6-TMPRSS2 cells, 2018/T424I showed similar growth kinetics compared to 2015/GD01, but significant reduction of infectious virus titers was seen in 2019/LAK and 2019/SV at 24 and 48 hours post infection (hpi) (Fig. 3E). This result was concordant with the reduced plaque size observed with these two viruses (Fig. 3D). The 2019/LAK virus showed a reduction of 3.04 and 1.43 log_10_ TCID_50_/ml at 24 and 48 hpi, respectively, compared with reference virus 2015/GD01, whereas the 2019/SV showed a lesser reduction of 1.46 and 0.52 log_10_ TCID_50_/ml at 24 and 48 hpi, respectively. Similar comparative reductions virus growth kinetics were seen in Calu3 cells with 2019/LAK and 2019/SV, although the magnitude of reductions observed was smaller (Fig. 3F). Area-under-curve (AUC) analysis showed 2015/I529T, 2019/LAK and 2019/SV produced less infectious virus as compared with 2015/GD01, in both cell cultures. In AUC analysis, we did not observe significant differences in replication kinetics with 2023/B5-3 and 2023/B5-4, compared with 2015/GD01 in both cell types (Fig. 3G and H). Reduction in replication was seen with 2023/B5-4 in VeroE6-TMPRSS2 cells at 48 hpi (0.49 log_10_ TCID_50_/ml) and with 2023/B5-3 in Calu3 cells at 72 hpi respectively (0.31 log_10_ TCID_50_/ml). Overall, these results suggest spike mutations from 2019 sequences could cause a reduction in viral replication, whereas mutations from 2023 sequences did not.

### Mechanisms of virus entry

We next investigated if any of these spike mutations impact the virus entry pathways of MERS-CoV, specifically the relative role of the cell surface mediated TMPRSS2 entry pathway and the endosomal cathepsin L entry pathway. In SARS-CoV-2, Omicron variant BA.1 showed a switch to the use of the endosomal pathway for cell entry, compared to the ancestral strains which predominantly used the cell surface pathway for cell entry [25]. In the case of Omicron BA.1, this entry pathway switch was contributed by mutations in the S2 domain. We observed several S2 mutations among our studied MERS-CoV spike sequences: Q914K in 2019/LAK, G705S and L1301V in 2019/SV and 3 conserved S743I, 1179-1180 del among the 2023/B5-1 to B5-4 spikes. We investigated the cell entry pathway preference using inhibitors targeting either TMPRSS2 (Camostat) or Cathepsin L (E64d) in VeroE6-TMPRSS2 cells, which offer both pathways for cell entry. Using recombinant viruses to infect cells pre-treated with entry inhibitors, we observed that only Camostat effectively inhibited viral replication, starting at 10 μM concentration (Fig. 4A). No effective inhibition was observed using the endosomal inhibitor E64d. Similarly, pseudoviruses expressing different MERS-CoV spikes showed the same entry preference via the TMPRSS2 entry pathway in VeroE6-TMPRSS2 cells (Fig. 4B). The data suggest that the clade B MERS-CoVs with spike mutations investigated here predominately utilize TMPRSS2 for cell entry.

**Figure 4.**
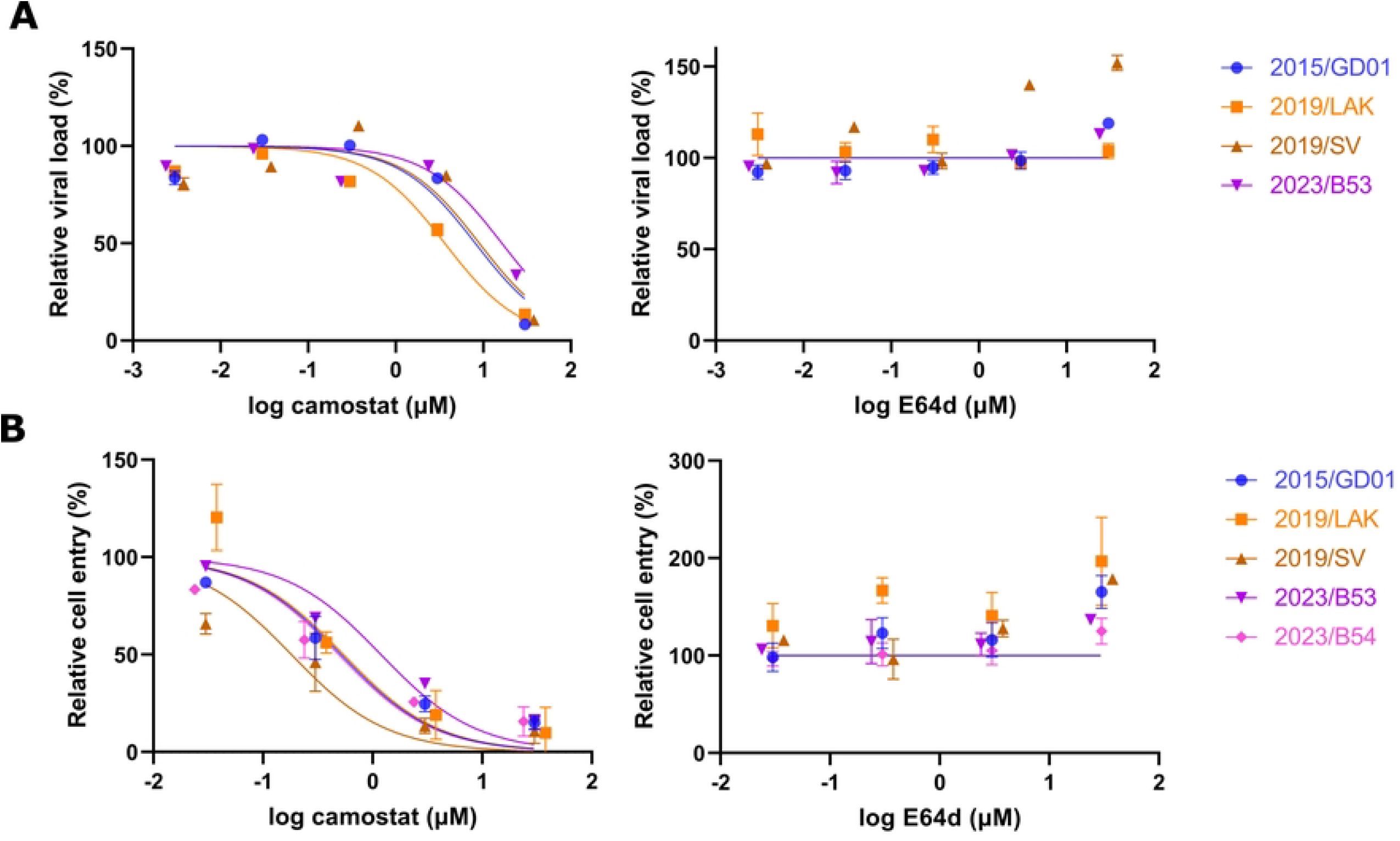
Entry pathways preference of MERS-CoV spike variants. Treatment of cells with camostat targeting the cell surface TMPRSS2 pathway and E64d targeting the endocytic cathepsin L pathway was used to block infectious virus entry via each pathway. Experiments with infectious virus (A) or pseudovirus (B) is shown for the different spike variants in VeroE6/TMPRSS2 cells. (A) Relative viral load was measured the RNA copies of MERS-CoV using the UpE assay in the cell supernatant at 48 hpi. (B) Relative cell entry was measured by RLU. (A-B) 2015/GD01 was the reference for comparison. Figures show the representative data from 2 independent experiments each with two biological replicates for each experiment each drug dose. An inhibition curve was fitted using the Hill equation with inhibition in Graphpad Prism.

## Discussion

Here, we assessed the phenotypic consequences of spike mutations that were observed to occur within the clade B lineage 5 MERS-CoV in relation to cell entry, viral replication and cell entry pathway preferences.

We developed a Golden Gate assembly platform that allowed robust rescue of recombinant MERS-CoV from full-MERS-CoV genome encoding pBAC infectious clones. Similar approaches have been applied from other SARS-CoV-2 studies [26, 27]. Based on the published sequences from Genbank (available from 2025 January), we synthesized the relevant fragment of the genome for our study. In about 30 days, the fragments were ready and available for the assembly reaction. With one additional week for plasmid preparation and two weeks of virus rescue experiments in a BSL3 lab, recombinant virus stock can be available for experiments in roughly 50 days for phenotyping experiments. A full MERS-CoV genome can be synthesized accordingly by partitioning the genomic sequence into separate plasmids (eg. pUC-F1 to F10 in this study). The Golden Gate assembly reaction is robust to generate the intact pBAC infectious clone with a high success rate. We believe this technique will be a valuable tool for MERS-CoV research to generate robust virus phenotyping data from new surveillance data and response the emerging threats from the ongoing evolution of MERS-CoV.

We showed that two types of spike mutations that occurred in 2019 MERS-CoV sequences, 2019/LAK and 2019/SV, did contribute to reduced cell entry and virus replication compared to an early clade B, lineage 5 virus 2015/GD01. On the other hand, although more recent 2023 viruses had acquired 8 to 10 spike amino acid substitutions in spike protein, they showed no significant difference in cell entry and viral replication compared with the ancestral clade B lineage 5 strain 2015/GD01. While our study highlighted that spike mutations in clade B MERS-CoV can be one factor determining the replication fitness of MERS-CoV, potentially impacting reduced zoonotic disease associated with viruses circulating in 2019, this was not true of more recent MERS-CoV circulating in 2023. The spike mutations did not appear to affect virus replication competence of the 2023 MERS-CoV. It is important to note that mutations parts of the genome other than spike protein may significantly impact virus replication competence and this deserves further study.

We identified a RBD V530A amino acid substitution and a transmembrane domain L1301V amino acid substitution that can contribute to a reduced cell entry in the 2019 spike sequences. The V530A mutation is one codon position adjacent to the I529T mutation, which has previously identified to be associated with reduced cell entry, virus replication and pathogenicity in mice [20, 21]. The I529T mutation was reported to have emerged later in the course of human-to-human transmission chains that occurred during the outbreak in the Republic of Korea in 2015 [28]. Further testing on the binding affinity of the V530A mutation can illustrate whether the spike codon 529-530 region is a hotspot for the virus to adapt in human via reduced DPP4 binding. For 2019/SV, the L1301V mutation located at the transmembrane domain of the spike was responsible for the reduce cell entry. The spike transmembrane domain consists of a hydrophobic domain, which is important for stabilizing the trimeric structure and membrane fusion. Replacing hydrophobic residues into hydrophilic residues in the transmembrane domain showed reduced psuedovirus cell-entry for SARS-CoV spike protein [29]. The leucine to valine residue change may possibly affect the hydrophobicity due to a larger isobutyl group in leucine compared to the isopropyl group in valine. Further experiments using a cell-cell fusion assay may elucidate the mechanism of the L1301V mutation.

While both V530A and L1301V spike mutation are associated with reduced cell entry and viral fitness, it is relevant to note that other human adaptive mutations elsewhere in the genome (e.g. nsp6 L232F) that increase viral fitness in human cells have been reported, and these 2015/GD01 viruses do in fact carry this nsp6 L232F amino acid substitution [30]. It is interesting to note that the extensive NTD and RBD mutations observed in 2023 sequences were not impacting the cell entry and viral replication in human cells, even though these mutations were located at the spike protein surface, in the respective domain. The V530A and L1301V amino acid substitutions observed in the 2019 MERS-CoV were no longer seen in the 2023 viruses. It is important to note that many of these mutations were driven by selection within the camelid host, rather than within humans, and thus, the phenotypic consequences of these mutations may or may not be functionally relevant in human cells. The impact, if any, of these mutations requires further experiments to be done in camelid cells or respiratory tissues.

Our studies indicate that the clade B lineage 5 viruses consistently prefer the use of TMPRSS2 cell-entry pathway. While DPP4 is more highly expressed in the epithelial cells of the bronchi, bronchioles, and alveoli, TMPRSS2 is expressed in both upper nasal epithelium and lower bronchi respiratory tracts [31, 32]. Ciliated epithelial cells, club cells, and alveolar type II pneumocytes have been shown to express TMPRSS2 and are susceptible to MERS-CoV infection [33, 34]. Our data showed that 2019 MERS-CoV viruses with reduced cell entry continued to use the TMPRSS2 entry pathway, that pathway of cell entry was not a reason for the reduced virus replication competence.

This study has a number of limitations. Our investigation was limited to the role of the spike protein. Although spike protein is well recognized as an important determinant of virus tropism and replication competence, which was also demonstrated with the 2019 viruses in our study, other regions of the genome can also contribute to viral fitness. Thus, we cannot exclude the possibility that reduced zoonotic cases in recent years may indeed be caused by viral genetic changes elsewhere in the genome. The 2023 viruses need to be isolated and phenotyped. The paucity of MERS-CoV sequence data in recent years also hinders a systematic analysis of the evolution and phenotype of these viruses. A second limitation is that our experiments were carried out in cell lines. Further experiments in ex vivo human respiratory tissues would be desirable to extend these findings.

In conclusion, spike mutations in recent 2023 sequences, though extensive, are not associated with altered virus replication competence and probably do not explain the reduced numbers of MERS cases observed in recent years. However, MERS-CoV in 2019 do appear to have reduced virus fitness as a consequence of spike mutations. Other potential explanations for the reduction in human MERS cases also need to be explore and these include the possibility of cross-immunity from COVID-19 infection or vaccination. Although COVID-19 infected or vaccinated individuals have no cross-neutralizing antibody to MERS-CoV, we had previously reported evidence that SARS-CoV-2 infection or vaccination does elicit spike S2 domain binding antibodies [35]. Such antibodies may contribute to some cross-immunity via antibody-dependent cell cytotoxicity or other functional pathways. Separately, cross-reactive T cell immunity may also potentially provide some cross-protection. These possibilities deserve investigation. Our study highlights that MERS-CoV continues to evolve with unpredictable consequences for pandemic risk. Intensive genetic and phenotypic surveillance of MERS-CoV is essential to understand the ongoing changes of the zoonotic potential of these viruses.

## Methods

### Phylogenetic analysis and spike mutation analysis

We analyzed 607 MERS-CoV genomes from Genbank with complete spike gene sequences excluding those arising for cell cultured isolates. The dataset was accessed on 2025 Jan7 and included recent sequences published by Hassan et al [19]. Sequences were aligned by MAFFT and subsequently analyzed to construct a PhyML tree using IQ-tree v2.1.4 using the standard substitution model selection [36]. Spike amino acid mutations against the MERS-CoV 2015/GD01 (accession number: KT006149.2) were counted from the alignment of spike protein sequences using MEGA v10.2.

### Pseudovirus production and infection

HEK293T (ATCC) cells were transfected with a plasmid mix of 850 ng of a pcDNA3.1 expression vector encoding a human codon-optimized MERS-CoV spike protein, 2.5 μg a lentiviral vector backbone encoding firefly luciferase and ZsGreen (Addgene plasmid #164432), 550 ng of a pHDM vector encoding HIV-1 GagPol (Addgene plasmid #164441), 550 ng of a pRC vector encoding HIV-1 Rev protein (Addgene plasmid #164443)and 550 ng of a pHDM vector expressing HIV-1 Tat protein (Addgene plasmid #164442) per 2 × 10^5^ cells using Trans-IT as transfection reagent. After 48 hours post-transfection, cell supernatant containing pseudoviruses was spun to remove dead cells for 5 mins at 1000g and was aliquoted and store at 80°C until use.

Pseudoviruses were quantified using a RT-qPCR strategy as described by Zhang et al with some modifications [37]. Pseudovirus-containing supernatants were RNA extracted and the RNAs were digested with DNase I according to the manufacturer’s protocol to remove any leftover plasmids. DNase-digested RNAs were reverse transcribed using Takara Perfect RT kit. Genomic copies of pseudoviruses were measured by a SYBR-green qPCR reaction (Applied Biosystems™) using a pair of primers targeting the luciferase gene (Fwd: 5’-CTTCGAGGAGGAGCTATTCTTG-3’ and Rev: 5’-GTCGTACTTGTCGATGAGAGTG-3’) and calculated from a standard curve with known copies of the lentiviral vector plasmid.

To infect target cells, 1 × 10^8^ genomic copies per 100 μl of pseudovirus was added to pre-seeded Vero and Calu3 cells incubated with 10% DMEM medium. After 48h of incubation in a CO_2_ incubator at 37°C, cell supernatants were removed and 30 μl of 1x lysis buffer were added for cell lysis for 5 mins. 100 μl of luciferase substrate (Promega E1500) were added to lysed cells and the luciferase activity was measured using the Perkin Elmer MicroBeta 2 instrument.

### MERS-CoV reverse genetics using pBAC infectious clones

To generate recombinant viruses with different spike variants, we utilized the Golden Gate assembly strategy to induce the desired mutations into a pBAC infectious clone encoding the MERS-CoV 2015/GD01 (obtained from Prof. Jincun Zhao, Guangzhou Medical University). We divided the MERS-CoV genomic region into 10 consecutive fragments and cloned each fragment separately into a pUC high copy number vector. Since the MERS-CoV genome contains 4 internal BsaI cut-site, synonymous mutations were induced into the fragments to abolish these cutsites for the assembly reaction (2438 nt C>A in F1, 8144nt C>A, 11681nt C>A and 11687nt A>G in F4, 21695nt C>T in F8 and 29384nt C>T in F10 respectively). Spike mutations were either induced into the pUC-F8 plasmid, that encodes a full-length spike gene using a site-directed mutagenesis kit (Agilent) or via gene synthesis (Genewiz). To prepare the assembly reaction, 75 ng of each plasmid (F1 to F10 plasmids, plus a pUC vector encoding the pBAC backbone) were added with 2 μl of T4 DNA Ligase Buffer (10X) and 2 μl of NEBridge Golden Gate Enzyme Mix (NEB). The reaction was put into a thermal reaction of 30 cycles of 37°C for 5 mins, followed by 16°C for 5 mins and ended with a step of 60°C for 5 mins. 1 μl of the reaction was electroporated into 25ul of NEB 10-beta Electrocompetent cells using the BTX ECM 630 and Bio-Rad GenePulser electroporators with a setting of 2.0 kV, 200 Omega, and 25 μF. Transformed cells were added with 950 μl of SOC medium and shaken for 37°C for 60 mins. 100 μl of cells were spread on chlorophenol LB agar plates to screen for positive clones. Colonies were picked and screened by PCR with primers flanking the cut-sites to confirm the ligation. pBAC from successful clones were extracted using NucleoBond Xtra Midi prep kit (Macherey-Nagel). Sequence identity of the pBACs were confirmed by NGS on iSeq 100 System (Illumina).

Rescue of recombinant MERS-CoVs from the infectious clones were performed as described previously [30]. Briefly, the pBAC was transfected to Vero cells using lipofectamine 2000 (ThermoFisher). 6 hours later, the transfection mix was removed and replaced with 2% FBS supplemented DMEM medium. After 72h post transfection, the supernatant was passage into VeroE6/TMPRSS2 cells and incubated for another 48h. The presence of CPE will indicate a successful rescue and the supernatant will be kept as P1 for a further next-round passage to generate the experimental stock. All virus stocks were sequenced by NGS, quantified for infectious viral titers using TCID_50_ assay in VeroE6/TMPRSS2 cells and aliquoted for storage at -80°C until use. Transfection of infectious clones and all experiments involved with rescued recombinant viruses were performed in a BSL3 facility in the University of Hong Kong.

### MERS-CoV infection and virus titration TCID_50_ assay

VeroE6/TMPRSS2 (kindly provided from Prof. Peter Cheung, Chinese University of Hong Kong) and Calu3 (ATCC) cells were seeded in 24-well plates one day prior with complete growth medium. Prior infection, cell medium was removed and washed with PBS once. Virus stocks diluted with 0% FBS DMEM medium to a dose of MOI 0.01 were incubated with cells for 1 hour at 37°C. After the incubation, virus inoculum was removed and cells were replenished with 2% FBS DMEM medium. Supernatants were harvested at 1, 24, 48 and 72hpi for virus titration. TCID_50_ assay was performed in VeroE6/TMPRSS2 cells. Cytopathic effects (CPE) from infected cells were measured at day 5 post infection.

### MERS-CoV spike entry pathway analysis using drug inhibitors

Entry inhibitors Camostat (Tocris, Cat.No. 3193) and E64d (Tocris, Cat. No. 4545) were serially diluted with 2% FBS DMEM medium and incubated on to VeroE6/TMPRSS2 cells for 2 hours prior infection. For live virus infection, a virus dose of MOI 0.01 was added to cells on top of the drug-containing medium. Supernatant was harvested at 48hpi and extracted for viral RNA for qPCR quantification using an UpE assay. For pseudovirus infection, 1 × 10^8^ genomic copies per 100 μl of pseudoviruses were added to drug-treated cells and the RLU was measured at 48hpi.

### Biosafety assessment

We completed an independent risk assessment of the genetic modification of MERS-CoV in this study in the Safety Office, the University of Hong Kong, prior the start of any virus rescue experiments. The spike mutations of 2015/I529T, 2019/LAK, 2015/SV, 2018/T424I, 2023/B5-4 and 2023/B5-3 introduced into the spike protein of MERS-CoV 2015/GD01 and rescued as infectious virus close were all previously reported to be present in MERS-CoV in the field and thus not novel mutations. Furthermore, each of these spike mutants was previously assessed in this study for virus entry using virus pseudotype experiments which demonstrated that the mutants did not exhibit any increase in virus entry, providing further assurance that these mutations were not likely to enhance virus replication. All laboratory studies with infectious MERS-CoV was carried out in the bio-safety level 3 laboratory at The University of Hong Kong. Virus psedotype experiments were carried out in bio-safety level 2 laboratory containment, as these are not capble of sustained virus replication.

## Acknowledgments

The research was funded by research grants from the US National Institutes of Health (contract no. U01-grant AI151810). R.T.Y.S. and M.P. conceived and planned the overall study, L.L.M.P. and Z.Z. provided advise on experimental design and KKML assisted with experiments. R.T.Y.S. wrote the original draft and all authors critically reviewed and revised the manuscript.

